# Interpreting Attention Mechanisms in Genomic Transformer Models: A Framework for Biological Insights

**DOI:** 10.1101/2025.06.26.661544

**Authors:** Micaela E. Consens, Ander Diaz-Navarro, Vivian Chu, Lincoln Stein, Housheng Hansen He, Alan Moses, Bo Wang

## Abstract

Transformer models have shown strong performance on biological sequence prediction tasks, but the interpretability of their internal mechanisms remains underexplored. Given their application in biomedical research, understanding the mechanisms behind these models’ predictions is crucial for their widespread adoption. We introduce a method to interpret attention heads in genomic transformers by correlating per-token attention scores with curated biological annotations, and we use GPT-4 to summarize each head’s focus. Applying this to DNABERT, Nucleotide Transformer, and scGPT, we find that attention heads learn biologically meaningful associations during self-supervised pre-training and that these associations shift with fine-tuning. We show that interpretability varies with tokenization scheme, and that context-dependence plays a key role in head behaviour. Through ablation, we demonstrate that heads strongly associated with biological features are more important for task performance than uninformative heads in the same layers. In DNABERT trained for TATA promoter prediction, we observe heads with positive and negative associations reflecting positive and negative learning dynamics. Our results offer a framework to trace how biological features are learned from random initialization to pre-training to fine-tuning, enabling insight into how genomic foundation models represent nucleotides, genes, and cells.

## Main

Recently, transformer models, originally designed for domains such as natural language processing (NLP)^1^ and computer vision^2^, have found success in genomic applications such as predicting regulatory elements in DNA^3,4^, correcting batch effects from scRNAseq data^5,6^, and more^7^.

Despite their success, transformer models, like most deep learning models, offer limited insight into their prediction mechanisms^8,9^. This lack of interpretability is particularly problematic in computational biology, where understanding the biological basis of predictions is crucial for scientific discovery and downstream clinical applications. While some work has explored interpretability in genomic deep learning^8,10–12^, these efforts often focus on downstream performance rather than what is learned during pre-training or fine-tuning.

A core component of transformer models is the attention mechanism, which allows models to prioritize different parts of the input sequence^13^. In genomics, this enables focus on relevant regions or genes, and because attention scores reflect this focus, they have been proposed as an interpretability signal^3,14^.

While attention has been examined in prokaryotic genome models^14^, and extensively in NLP and protein models^15–19^, most human genome and single-cell transformer models lack systematic analysis of their attention heads. Furthermore, with over 100 attention heads, manual interpretation of these models is impractical.

We propose a scalable method to automatically assign biological interpretations to each head. We apply this method to three models: DNABERT^3^ (pre-trained DNA model), Nucleotide Transformer^4^ (pre-trained DNA model), and scGPT^5^ (pre-trained single-cell model), spanning DNA and single-cell tasks. We show that many heads are associated with biological annotations, and these preferentially contribute to predictive performance. We find that associations with biological annotations arise during pre-training, and that head-wise attention-feature correlations can be summarized by GPT-4 to yield biological meaning^20,21^. This provides a new framework to evaluate what genomic transformers learn during pre-training and fine-tuning.

## Results

### Interpreting attention heads in genomic transformer models

This study focuses on three models, each trained for two distinct tasks, to demonstrate the generalizability of our method across various transformer architectures and datasets.

DNABERT and Nucleotide Transformer are pre-trained language model-style transformers that can be fine-tuned for different tasks such as enhancer or TATA/non-TATA promoter prediction. DNABERT processes DNA sequences using a 510-nucleotide window, while Nucleotide Transformer (specifically, nucleotide-transformer-v2-500m-multi-species) processes sequences of up to 6,000 nucleotides through non-overlapping 6-mer tokenization. In contrast, scGPT is a transformer model trained on single-cell gene expression data, which we fine-tuned on two datasets for cell type classification.

An overview of the interpretability workflow is shown in Figure 1. We applied the full analysis pipeline to each model (DNABERT, Nucleotide Transformer, and scGPT) in three forms: randomly initialized, pre-trained, and fine-tuned. We also evaluated a variant of DNABERT pre-trained on random k-mers, as described by Zhang et al.^22^

**Figure 1.**
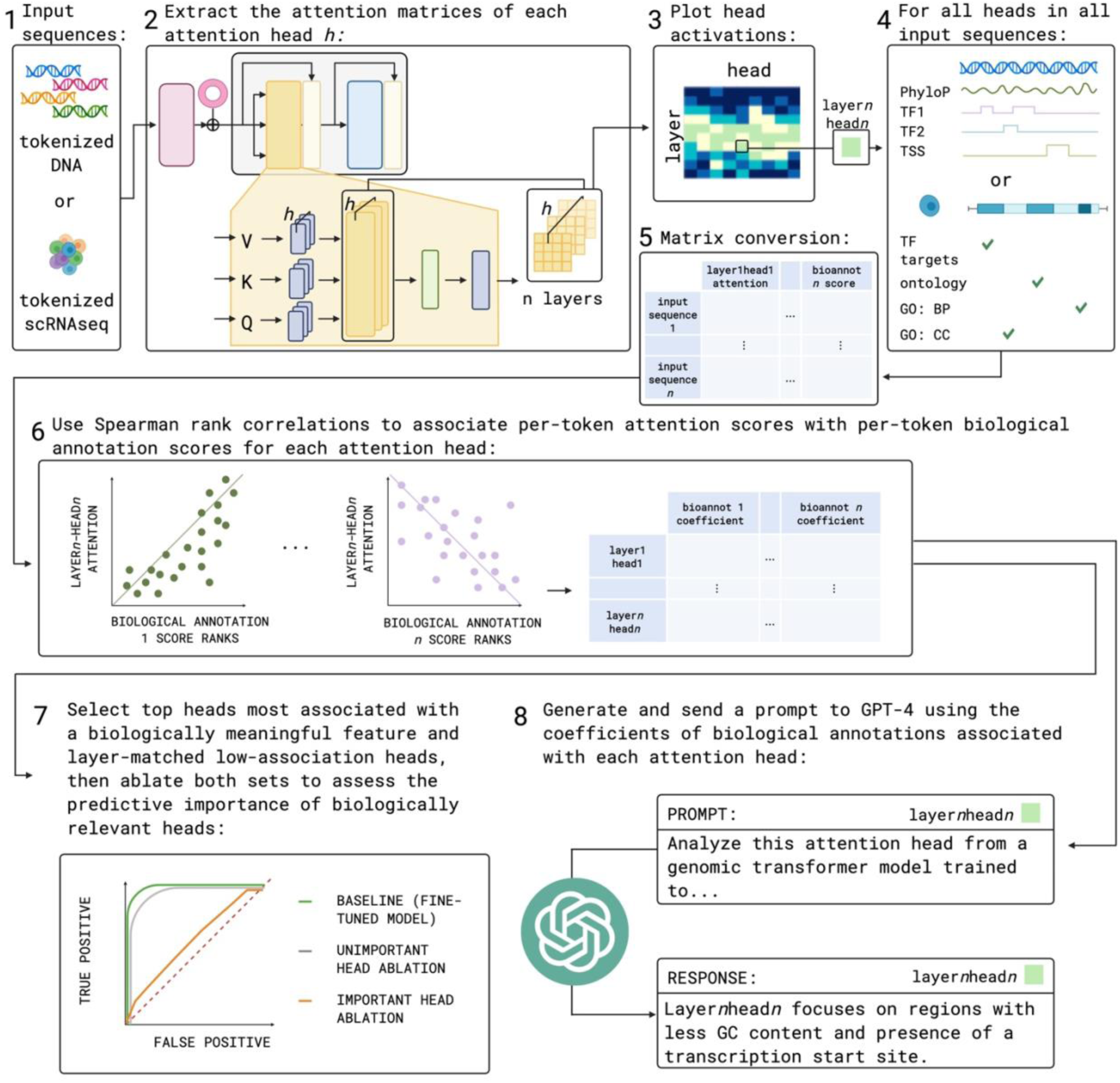
Flowchart illustrating the main steps of the genomic interpretability analysis. 1) DNA sequences (DNABERT, Nucleotide Transformer) and per-cell gene counts (scGPT) are tokenized for input to the models. 2) Attention scores are extracted from all layers and heads for each token of the individual input sequences/cells for a randomly initialized version of each models’ architecture, and then for both the pre-trained and fine-tuned versions. 3) Model mean head activations are plotted to visualize focus areas. 4) biological annotations are then mapped to the input tokens and 5) merged with their per-token attention scores. 6) These features are then associated with the attention scores using Spearman correlation. For each label in the classification task, the feature correlations are calculated only on the subset of those samples to get label-specific head correlations. 7) The top associated heads for a given feature in the finetuned models are identified and selected, along with layer-matched low-association heads. These heads are then both ablated and the model’s performance is assessed to determine whether biologically interpretable heads are more, less, or of the same importance as non-interpretable heads. 8) Using the coefficients derived from the attention-feature annotation correlations, a prompt is generated and sent to GPT-4 to zero-shot summarize each attention head’s activity in all models but the randomly initialized baselines.

For each model, we extracted the attention scores of every head in every layer using curated evaluation datasets (*see Methods*). These scores were compiled into matrices for each model containing attention scores per token and corresponding biological annotations (Supplementary Table S1). For DNABERT and Nucleotide Transformer, we focused on genome-mappable features such as transcription start sites (TSS), transcription factor (TF) binding sites, repeats, phyloP scores, and GC content. For scGPT, we annotated genes using pathway and gene ontology terms (*see Methods*).

We computed Spearman rank correlations for each attention head by comparing the per-token deviation from that head’s mean attention score to the corresponding biological annotations. Since Spearman correlation is invariant to monotonic transformations such as centering, this is equivalent to correlating raw attention scores with biological features. For fine-tuned models, we provided the correlation coefficients to GPT-4 in a zero-shot setting to assign descriptive summaries to each attention head. We also conducted ablation tests on the fine-tuned models, comparing the impact of ablating heads strongly associated with important biological features to that of ablating layer-matched heads with weak associations.

We found that the tokenization scheme influences model interpretability using our method. In a synthetic TATA identification task, removing DNABERT attention heads directly associated with the TATAAA motif had the greatest impact on performance. In contrast, for Nucleotide Transformer, removing heads associated with GC content – likely acting as proxies due to token fragmentation – had greater effect (*see Methods*).

### Attention heads in genomic transformer models are context dependent

A preliminary analysis of the raw attention scores on the k-mers alone revealed that attention scores in genomic transformer models are context dependent. Attention heads selectively assign higher scores to tokens related to certain biological features – such as the TATAAA k-mer in the DNABERT TATA model – when the entire sequence corresponds to a TATA promoter (Figure 2). This suggests that the transformer’s attention mechanism leverages other sequence features to determine whether the TATAAA k-mer is part of a promoter. We show that certain attention heads, such as layer10-head0, are specialized for detecting context-specific features, with context dependence reflected as a relative increase in attention compared to the head’s mean. To highlight these relative increases, we computed per-token deviations from the head’s mean attention score (Supplementary Figure S1). However, since our method uses Spearman correlations, which are invariant to shifts in mean, this approach yields results identical to correlating raw attention scores with biological features.

**Figure 2.**
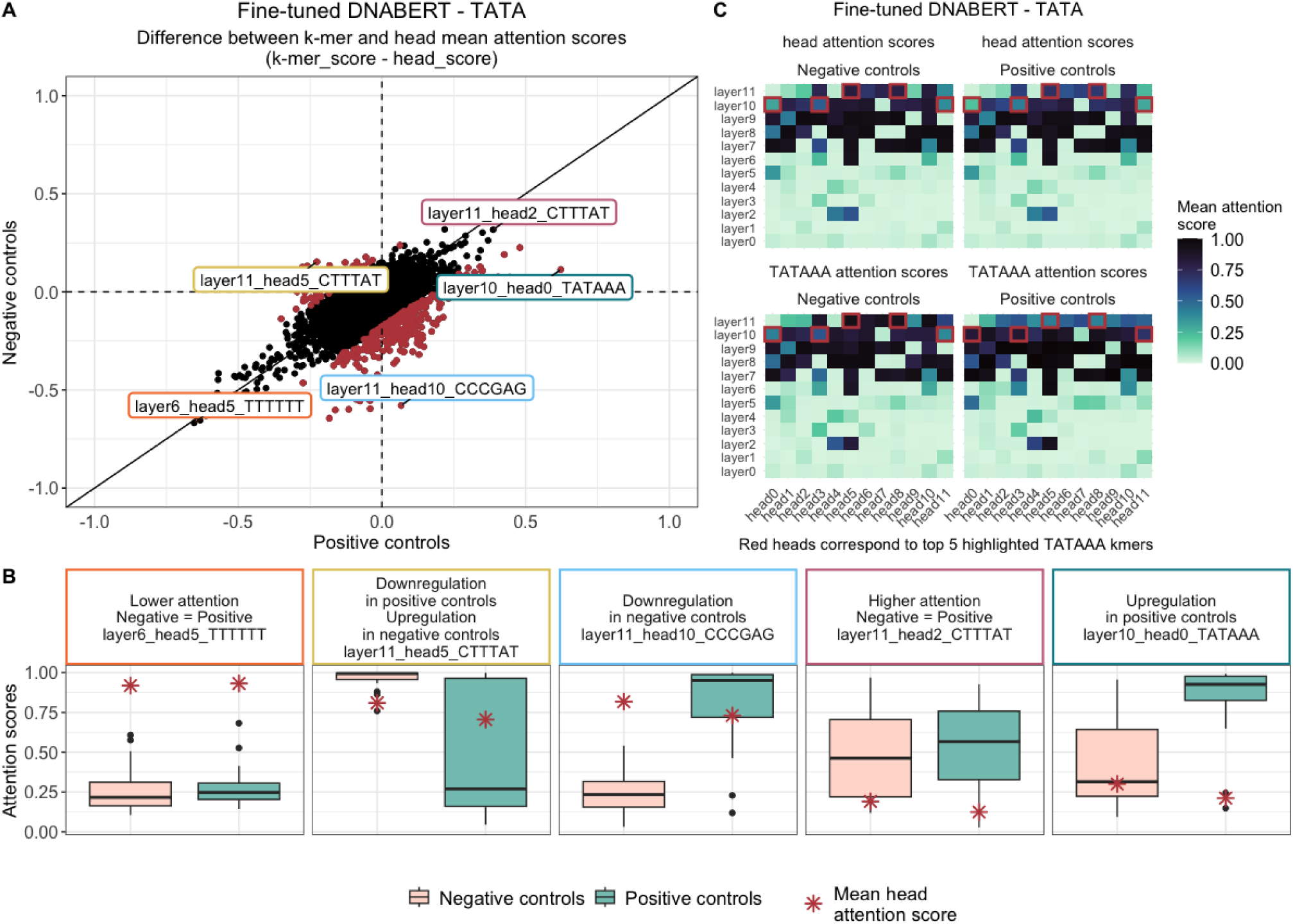
Visualization of differences in k-mer attention scores depending on sequence context for the DNABERT model fine-tuned on the TATA promoter prediction task. A) Scatter plot showing the difference between the mean head attention score and the specific k-mer mean attention score, with sequences labeled as promoters (positive controls, x-axis) or non-promoters (negative controls, y-axis). Red-highlighted k-mers represent values beyond three standard deviations. B) Boxplots displaying the distribution of specific k-mer attention scores, helping to interpret the patterns seen in the scatter plot. C) Heatmap showing mean head attention and TATAAA k-mer mean attention across all heads and layers, separated by promoter (positive) and non-promoter (negative) sequences. Highlighted heads indicate the top five with the strongest differences between sequence types and between global vs. k-mer mean attention.

### Genomic transformer model attention heads learn associations with biological features during pre-training – even random pre-training

We quantified the associations between attention scores and biological features using Spearman correlation, reporting z-scores derived from label-preserving random shuffles. To assess what is learned during pre-training, we compared z-scores in pre-trained models to those from randomly initialized versions (Figure 3A and 3D and Supplementary Figure S2). This comparison was performed separately for each label class.

**Figure 3.**
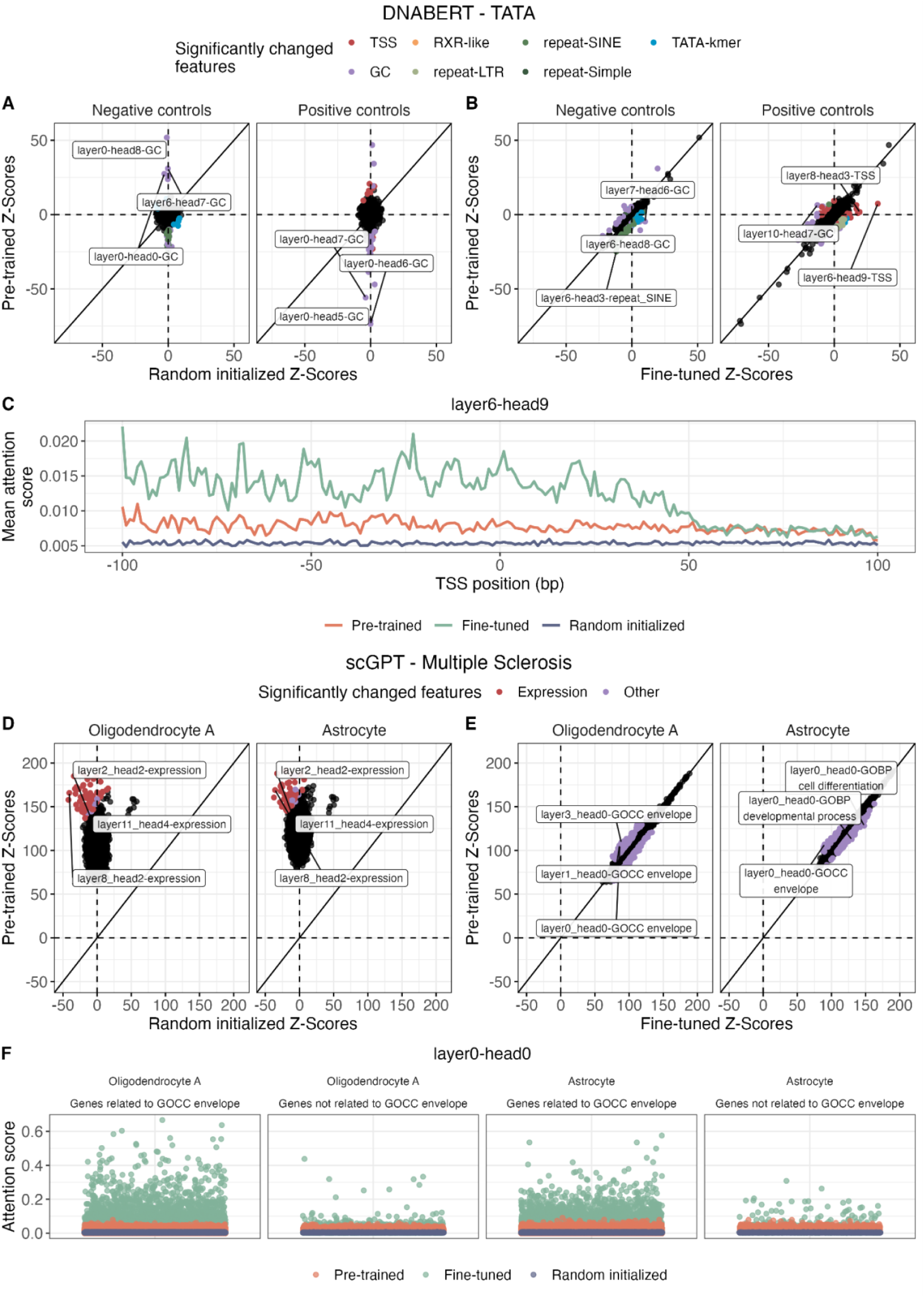
Visualization of feature importance in randomly initialized, pre-trained, and fine-tuned DNABERT (TATA promoter prediction task) and scGPT (Multiple Sclerosis dataset) models. A–B) Z-scores for each feature and attention head in the DNABERT models: A) comparison between randomly initialized and pre-trained models; B) comparison between pre-trained and fine-tuned models. Scores are shown separately for sequences that are real promoters (positive controls) and non-promoters (negative controls). C) Mean attention score across 200 bases surrounding the transcription start site (TSS) for head 9 in layer 6 –identified as the feature with the largest difference between the pre-trained and fine-tuned models in positive control sequences. D–E) Z-scores for each feature and attention head in the scGPT models: D) comparison between randomly initialized and pre-trained models; E) comparison between pre-trained and fine-tuned models. Since the Multiple Sclerosis dataset is a multiclass classification task, the two cell types with the largest number of cells were selected (Oligodendrocyte A, n = 154; Astrocyte, n = 107). F) Attention score of each gene for head 0 in layer 0, depending on whether the gene belongs to the “Envelope” cellular component GO term, shown separately for Oligodendrocyte A and Astrocyte cells. For A-B and D-E scatterplots, features that deviate more than three standard deviations from the diagonal are considered significant. The top three such features are labelled.

In the pre-trained DNABERT TATA model, several attention heads show strong associations with known biological features (e.g., layer0-head8 with GC content in negative controls, Figure 3A), whereas the randomly initialized model shows no significant associations. Similarly, in the pre-trained scGPT Multiple Sclerosis (MS) model, expression-related features received substantially higher attention compared to the randomly initialized version. This pattern was consistent across cell populations, suggesting that the pre-training effectively captures cell-type specific expression signatures relevant to MS pathology. Comparable patterns were observed across other models and tasks (Figure 3A and 3D and Supplementary Figure S2), indicating that pre-training consistently induces biologically meaningful attention patterns. This mirrors findings in NLP from Clark et al.^19^, where pre-trained attention heads were shown to learn syntactic information.

Interestingly, in the DNABERT model pre-trained on random DNA 6-mers from Zhang et al.^22^, we found that the z-score patterns more closely resemble those from the pre-trained model than from the randomly initialized one (Supplementary Figure S3). This suggests that even random DNA sequences can provide enough structure for models to learn generalizable biological signals, consistent with the performance of randomly pre-trained models reported by Zhang et al.

### Genomic transformer model attention head-feature associations change during fine-tuning

To assess how fine-tuning affects model attention patterns, we repeated the z-score analysis on all the fine-tuned models and compared the results to their pre-trained counterparts. Figures 3B and 3E illustrate these comparisons for the DNABERT TATA and scGPT MS models, respectively. Pre-training induces substantial shifts in feature associations, transforming many heads from exhibiting near-zero correlations to significant correlations with biological features. In contrast, fine-tuning primarily adjusts heads along the diagonal, suggesting that most heads retain their pre-trained associations (e.g., GC-related heads for the DNABERT TATA task). This trend is consistent across other models and tasks (Supplementary Figure S4). However, a subset of heads shift focus during fine-tuning to become more strongly associated with task-relevant features, such as the TSS for DNABERT TATA (Figure 3B) and GOCC (Gene Ontology Cellular Component) envelope features for scGPT fine-tuned on the pancreas dataset (Figure 3E). This is supported by findings from Kovaleva et al.^18^, who showed that attention patterns change from pre-training to fine-tuning in DNABERT models trained on natural language.

In the DNABERT TATA model, for instance, layer6-head9 shows a strong increase in correlation with the TSS after fine-tuning. Figure 3C <u>highlights a localized increase in attention near the TSS</u>, which tapers off beyond 50 bp downstream. While DNABERT’s original paper reported increased attention upstream of the TSS across all heads^3^, our analysis identifies the specific heads driving this pattern. Although the raw correlation is modest, the association is statistically significant as a z-score, indicating a consistent, if weak, pattern. This means the variation in layer6-head9’s attention cannot be explained solely using the TSS feature and could be influenced by a combination of signals such as local sequence context or co-occurring features. While we label this head as a TSS-specific head, we acknowledge this is an oversimplification of the attention head’s activity.

Similarly, in scGPT MS (Figure 3F<u>), fine-tuning enhances</u> attention specialization for different biological contexts. For example, layer0-head0 shows elevated attention scores for GOCC envelope-related genes compared to both pre-trained and randomly initialized models in oligodendrocyte A and astrocyte populations. The consistently positive feature associations suggest that fine-tuning accentuates, rather than redirects, the biological focus established during pre-training. Together, these results indicate that pre-training lays a foundation of general biological knowledge that fine-tuning refines for specific tasks.

### Removing attention heads associated with biological features impacts model performance more than layer-matched heads

Across all models and datasets, ablating attention heads with strong associations to biological features led to larger drops in performance compared to ablating layer-matched heads without such associations. This pattern held across both DNA and transcriptomic models, and across tasks ranging from simple motif detection (e.g., TATA promoters) to more complex classification (e.g., enhancer detection and single-cell domain labels) (Supplementary Figure S5). Representative examples are shown in Figure 4 for DNABERT (TSS), Nucleotide Transformer (TSS), and scGPT (GOCC-neuron projection).

**Figure 4.**
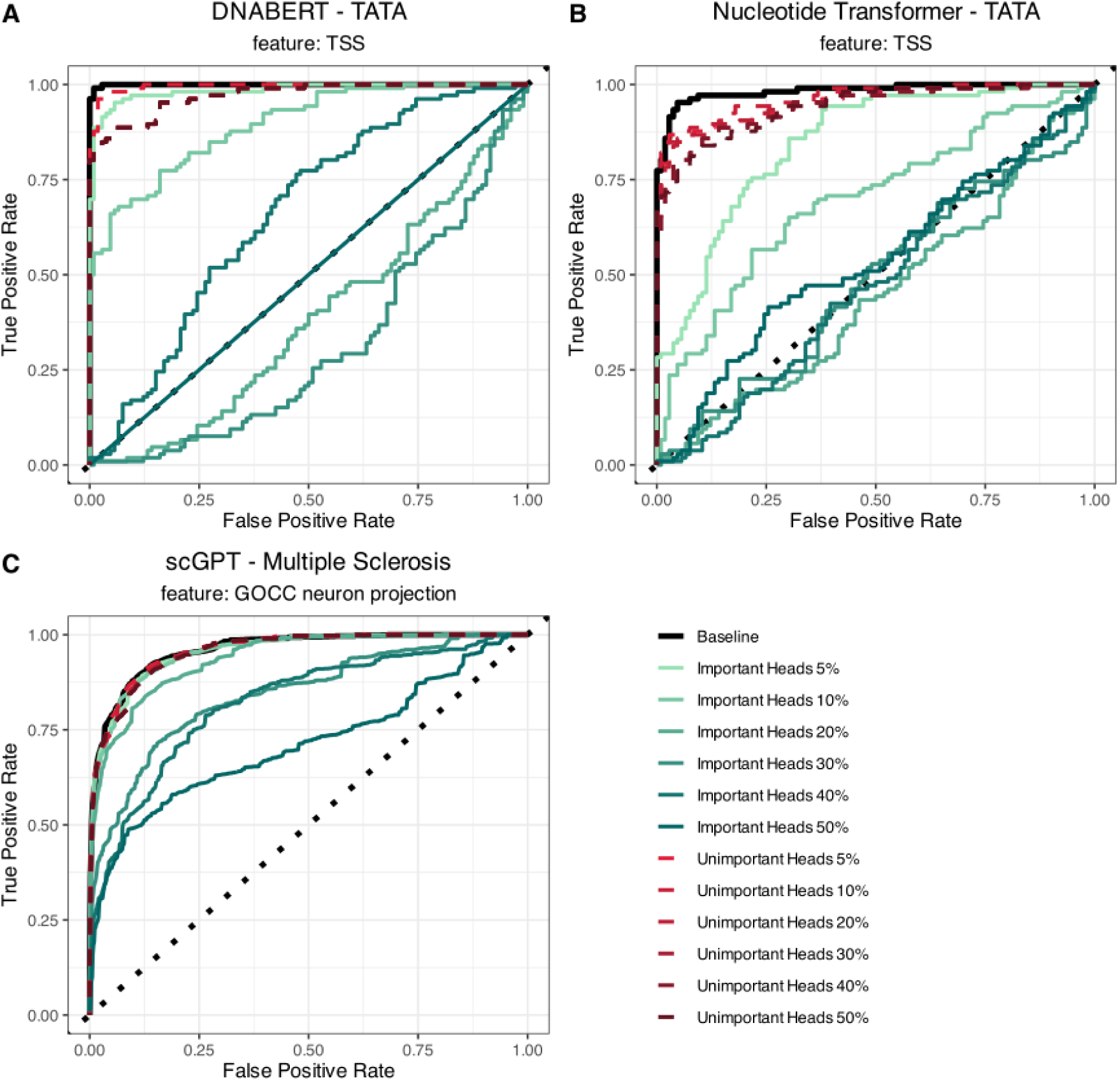
ROC curves for the head ablation experiments. A–B) DNABERT and Nucleotide Transformer models fine-tuned on the TATA promoter prediction task. The ablated heads are those correlated with the transcription start site (TSS) feature. C) scGPT fine-tuned on the Multiple Sclerosis dataset, with ablation of heads correlated with the “neuron projection” cellular component GO term. The dotted diagonal line represents a random classifier (AUC = 0.5).

These findings parallel prior work in NLP, where Voita et al.^23^ showed that important heads that remain after pruning tend to exhibit interpretable behaviors. While Voita et al.’s experiments relied on gradient-based identification of important heads, and our method relies on attention-feature correlations, both approaches highlight the predictive value of interpretable heads.

Aligning with Michel et al.^17^, who found that many heads (up to 40%) can be removed without affecting performance, we observed that substantial ablation (including of significantly-associated heads) is often required to meaningfully impact model accuracy. This indicates that, for certain tasks and features, the learned genomic information is distributed across many heads. Overall, our results suggest that, in genomic transformer models, biologically meaningful heads tend to be functionally important for prediction.

### In DNABERT fine-tuned to predict TATA-containing promoters we observe positively and negatively associated attention heads reflect positive and negative learning

In DNABERT TATA, ablating 20-30% of attention heads reduced performance below random chance, indicating prediction inversion (Figure 4A). This suggests the model misclassified positive and negatives more frequently than by chance, particularly when the ablation set included many heads strongly negatively associated with the TSS feature. A similar pattern was observed for the TATAAA k-mer feature (Supplementary Figure S6A).

We hypothesize that positively and negatively associated heads reflect two distinct modes of learning: *positive learning*, where heads attend more to predictive features (e.g. the TATA k-mer, or TSS), and *negative learning*, where heads suppress attention to those regions, effectively learning from feature absence.

To test this, we stratified attention heads by the direction of their correlation with the TATAAA k-mer and ablated the most positively or negatively associated heads. Removing the top positively associated heads caused AUC to decline gradually, with inversion occurring around the 40% ablation mark (Supplementary Figure S6B), similar to but slightly better than the association-direction agnostic ablations at 20-30%. This suggests the TATA k-mer is learned in a distributed manner, requiring many heads to be ablated before performance is substantially affected. In contrast, ablating a small fraction (5%) of the most negatively associated heads caused immediate performance degradation and decision boundary flipping (Supplementary Figure S6C). Further ablation of negatively heads had little additional effect, likely due to the small number of significantly negative heads and their small associations (Note: there were very few significantly negatively associated heads compared to positively associated heads).

However, attention may act in a context-specific manner. Globally negative heads may still increase attention under certain label conditions, and vice versa. This may explain why ablating only positively associated heads also led to performance inversion at higher thresholds, positive and negative associations are not strictly linear or label-agnostic. Thus, interpreting head behaviour solely based on global correlations risks oversimplifying how heads contribute to decisions.

Together, these results support a working model where attention heads can attend to either the presence or absence of features. Further validation is needed, ideally through label-specific ablations and context-aware interpretation strategies.

### GPT-4 can zero-shot summarize attention head activity, but explanations need to be verified

We formatted each attention head’s z-scores for all possible features in the fine-tuned models, such that we could send them zero-shot to GPT-4 to summarize. For DNABERT’s layer6-head9, GPT-4 returned an interpretation consistent with observed patterns in Figure 3C, describing the head as “TATA Box and Core Promoter Recognition.” GPT-4 returned an explanation of the head as “*This head strongly emphasizes the canonical TATAAA motif (8 vs –4) and transcription start site signals (33 vs 4), clearly distinguishing TATA-promoter sequences from non-promoters. It also highlights positive signals from phyloP conservation (6 vs 0) and bHLH and TF_bZIP motifs with positive values in TATA-promoters but negative or zero in non-promoters, reflecting known transcription factor binding and evolutionary conservation typical of functional core promoters*.” This explanation aligns with the z-scores, with GPT-4 effectively reporting that positive sequences are associated with the TATAAA motif with a z-score of 8, compared to negative sequences with a z-score of –4. GPT-4 also correctly reports the positive and negative z-scores for the TSS and phyloP as well (33 and 6 in positive, and 4 and 0 in negative). This suggests GPT-4 is able to correctly summarize and interpret the z-score information in a label-specific way, and even draw on prior biological knowledge to explain the correlations.

We also examined layer0-head0 for scGTP fine-tuned to predict MS, which is described by GPT-4 as a Neuronal Development and Synaptic Function head. GPT-4 explains the head-activity as “*This head captures gene expression programs related to neuron differentiation and synapse organization, with the strongest coefficients in excitatory neurons (especially cortical layers 2-6) and interneurons (including SST, PVALB, and VIP subtypes). The elevated values for neuron projection, synapse, and neurogenesis GO terms align with the distinctive transcriptional profiles of mature neurons, differentiating them from glial and immune cells in the CNS*.” GPT-4 does not highlight the GOCC feature we show in Figure 3F, but this may be because other terms have higher z-scores. This highlights that even the explanations given by GPT-4 overlook important associations when there are multiple label-specific correlations, and therefore are also a simplification of attention head activity.

While these summaries provide a way to navigate the multitude of z-scores for all the biological features, stratified in a label-specific manner, GPT-4 generated explanations may be over-simplified, and must be validated by systematic evaluation of the true z-scores and biological ground truth information^21^.

## Discussion

Overall, we aim to interpret the attention heads of two DNA transformer models and one single-cell transformer model (DNABERT, Nucleotide Transformer, and scGPT) in terms of biological annotations. We compute Spearman rank correlations between attention scores and annotation scores per token for each attention head, and apply GPT-4 to summarize head activity in biological terms. Ablation experiments evaluate the contribution of interpretable heads to task performance.

We show that attention heads are context-dependent, with behavior varying across class labels. This underscores the need for label-specific analyses, as global correlations may overlook heads that are informative only within subsets of samples.

Biological associations emerge during pre-training and are modulated by fine-tuning. Even the DNABERT model pre-trained on random sequences learned generalizable features during pre-training, supporting that structure in synthetic sequences is enough to drive learning.

Furthermore, many associations to biological features learned during pre-training are maintained or emphasized during fine-tuning. For a subset of heads, their attention associations shift substantially to be more task-specific, such as focusing on the TSS for TATA promoter prediction.

Tokenization affects interpretability. In a synthetic dataset, DNABERT’s overlapping k-mers made it easier to detect TATA motifs. In contrast, Nucleotide Transformer’s non-overlapping 6-mers diluted this signal but amplified GC-content associations. The GC associated heads could identify the TATA motif within sequences, even though positive and negative sequences across the training and test dataset were GC-balanced. This highlights that tokenization choices shape our ability to interpret what models’ learn and potentially changes the way these models learn.

Heads highly correlated with biological features are more important for prediction than uninformative, layer-matched heads. We also observed preliminary evidence for distinct positive and negative learning dynamics, heads that focus on either the presence or absence of features, though further work is needed to confirm this pattern.

Our goal is not to claim that attention heads operate solely on interpretable biological features, but to show that some head activity can be meaningfully explained in biological terms. We accept, as others have noted^21^, that these correlational mappings are imperfect and limited by human annotations.

Futhermore, we acknowledge limitations of attention as an interpretability tool. Attention weights can reflect relevant features but are unstable and layer-dependent^8^. Moreover, attention applies to tokens at the input layer, but we interpret scores across all layers as if token aligned. Gradient-based methods could better capture input relationships^24^. Despite this, attention has been previously used for interpretation^25,26^, and we build on this foundation.

Our curated biological features introduce interpretive bias. Correlations are constrained to selected annotations, whose quality and resolution vary. Model inference data also affects interpretability. Furthermore, we simplify context-dependence to class labels, overlooking intra-class heterogeneity (e.g., tissue-specific enhancers). Many attention-feature correlations are weak, possibly due to unmodeled combinatorial interactions. Models may also focus on features not captured in our annotations, and GPT-4 may overgeneralize in its summaries.

Future work should refine annotation selection, account for inter-token dependence, and model interactions between features to reveal more complex attention patterns. Given the limitations of attention, other methods like Layer-wise Relevance Propagation (LRP) should be used to trace relevance across model layers^24,27^. Despite these limitations, our findings offer a practical framework for interpreting attention in genomic foundation models, particularly for looking at the role of many heads at once, and examining what these models learn from random initialization, to pre-training, to fine-tuning.

## Methods

Our methodology involves both direct analysis of attention heads and interpretation assisted by GPT-4. For each model, we first collected all of the attention scores for every head and layer using a curated dataset with biological annotations of interest annotated per token. We then conducted a detailed analysis of a subset of these heads to determine their biological relevance. For the remaining heads, we utilized GPT-4’s prompting capabilities to generate hypotheses about their potential biological roles. Our approach is based on OpenAI’s previous work explaining neurons in large language models (LLMs)^21^, but instead of explaining every neuron in an NLP model, we explain every attention head in a DNA or single-cell transformer model.

### Model summary and experimental setup

#### DNABERT

DNABERT is a language model for genomics that is built off of the original BERT architecture. The model has 12 attention heads in each of its 12 transformer layers. During pre-training, DNABERT is trained using Masked Language Modelling (MLM) to predict masked k-mer (k=6) tokens from the entire human genome. During fine-tuning, the pre-trained weights are loaded into a BERT model for classification, where no weights are frozen from the pre-training, and additional weights for classification layers must be learned in fine-tuning. The authors of DNABERT released the pre-trained weights of the model at the time of publishing, and these weights can be fine-tuned for a variety of downstream tasks including classifying proximal and core promoter regions or identifying enhancer sequences from non-enhancer sequences.

We trained the DNABERT model for a total of three tasks. One of these tasks was for predicting enhancer sequences, another was for predicting TATA promoters from non-promoter sequences (a biologically straight-forward task), and a third was to predict whether randomly generated DNA sequences contained a TATAAA box within (as a control). For simplicity, we refer to these tasks as enhancer prediction, TATA prediction, and fake TATA prediction. The models were fine-tuned on a single NVIDIA A40 GPU with 48GB memory using standard settings in the fine-tuning pipeline on the DNABERT GitHub (Table 1).

**Table 1.**
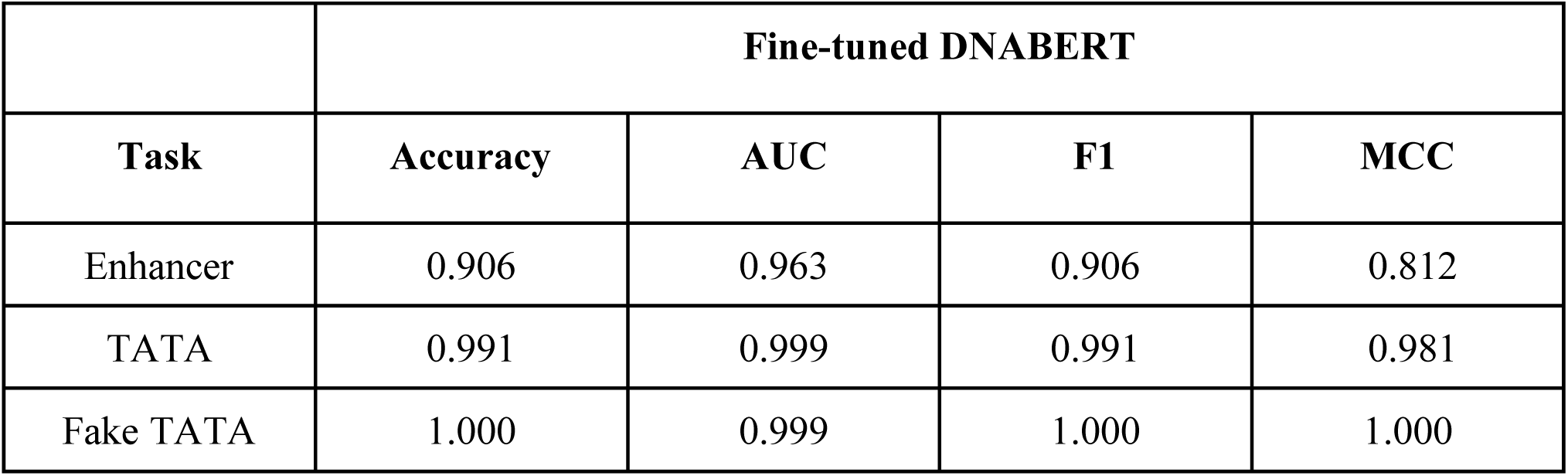
DNABERT Model Performance Fine-tuned.

For DNABERT, we additionally examine a version of the model pre-trained on random k-mers, instead of on k-mers from the human genome. This model is from the Zhang et al. paper^22^ that investigated DNABERTs k-merization and pre-training, and found a DNABERT pre-trained on random k-mers had similar performance on downstream tasks to a DNABERT pre-trained on the human genome. We download the DNABERT model pre-trained on random k-mers with k=6 from https://github.com/yaozhong/bert_investigation/blob/main/pt_models/download_url.txt.

#### Nucleotide Transformer

The Nucleotide Transformer is a language model for genomics that follows the BERT architecture. Like DNABERT, it was trained using MLM, but on data from 850 species rather than only humans. The model accepts inputs of up to 1,000 tokens (equivalent to 6,000 nucleotides based on non-overlapping 6-mer tokenization) and consists of 29 layers, each with 16 attention heads. We fine-tuned the model available on HuggingFace at https://huggingface.co/InstaDeepAI/nucleotide-transformer-v2-500m-multi-species on the same three tasks that DNABERT was trained on, on a single NVIDIA RTX A6000 GPU with 48GB of memory using standard settings from the Nucleotide Transformer HuggingFace. (Table 2).

**Table 2.** Nucleotide Transformer Model Performance Fine-tuned.

|  | **Fine-tuned NT** |  |  |  |
| --- | --- | --- | --- | --- |
| **Task** | **Accuracy** | **AUC** | **F1** | **MCC** |
| Enhancer | 0.8730 | 0.9455 | 0.8685 | 0.7477 |
| TATA | 0.9481 | 0.9834 | 0.9484 | 0.8963 |
| Fake TATA | 0.9940 | 1.0000 | 0.9940 | 0.9881 |

#### scGPT

scGPT is a large-language model for genomics that takes as input tokenized single-cell sequencing data (scRNA-seq, scATAC-seq), where each token represents a gene identifier. For scRNA-seq data, the token is associated with gene expression levels, while for scATAC-seq data, it corresponds to chromatin accessibility in a peak region. In scGPT, each cell is analogous to a ‘sentence’ composed of genes. scGPT can be fine-tuned for many downstream tasks, including cell-type classification, batch correction, and multi-omic integration. The model consists of 12 stacked transformer layers with 8 attention heads each, and can either run Flash Attention^28^ or full attention on the inputted sequences. For this paper, we implemented scGPT with full attention in order to retrieve the attention scores of every head and layer in the model.

The whole-human foundation model for scGPT was pre-trained using data from the CELLxGENE collection (https://cellxgene.cziscience.com/) using the Census API of 33 million normal human cells of 51 organs or tissues and 441 studies. The whole human pre-trained model was downloaded from the authors’ GitHub (https://github.com/bowang-lab/scGPT). The pre-trained scGPT was fine-tuned for cell type annotation, taking the transformer output cell embeddings as input and outputting categorical positions for cell types. For this project, we trained scGPT for two different tasks: classifying cell types from the multiple sclerosis (MS) dataset, and classifying cell types from the pancreas dataset (Table 3). The whole model was trained with cross-entropy on a reference dataset with expert annotations and then used to predict cell types on a held-out query data partition. We set a maximum input length of 500 for all tasks. For cells with a number of non-zero genes larger than the maximum input length, 500 input genes would be randomly sampled at each iteration, uniformly sampled from the three options of 0.25, 0.50 and 0.75.

**Table 3.** scGPT cell type annotation performance on MS and human pancreas datasets.

| Dataset | Accuracy | Precision | Recall | Macro F1 |
| --- | --- | --- | --- | --- |
| Multiple Sclerosis | 0.820 | 0.536 | 0.507 | 0.514 |
| Human Pancreas | 0.930 | 0.767 | 0.757 | 0.755 |

#### Random Initialization

For random initialization of models (DNABERT, Nucleotide Transformer, and scGPT), we employed a standardized approach as described in Vishniakov et al.^29^ using a normal distribution N(0, 0.02) across all model architectures. Specifically, we initialized linear layer weights from N(0, 0.02) with biases set to zero, embedding layer weights from N(0, 0.02), and LayerNorm parameters with scaling factors (gamma) set to 1.0 and bias terms (beta) set to 0.0.

### Datasets

#### Genome annotations

To associate the attention heads with different genomic features, we mapped each DNABERT and Nucleotide Transformer k-mer to experimentally validated transcription factor binding sites (TF), repeat elements and transcription start sites (TSS), using the TFLink, RepeatMasker and refTSS v4.1 databases^30–32^, respectively. In addition to the previous features, conservation scores across species were annotated using phyloP scores from the UCSC Genome Browser database.

#### Enhancer sequences

The datasets for identifying human enhancers were taken from a recent paper^33^ that uses the Ensembl v100 database^34^(p 202) and the VISTA Enhancer Browser^35^ (https://huggingface.co/datasets/katarinagresova/Genomic_Benchmarks_human_enhancers_ensembl). Due to the variable length sequences and ‘N’ encoded nucleotides in these datasets, we processed them by removing all sequences shorter than 50 bp or longer than 510 bp, as well as those containing ‘N’ nucleotides. After pre-processing, the training dataset consisted of 57,827 enhancer and 57,768 non-enhancer (control) sequences, whereas the test dataset contained 14,475 enhancer and 14,466 control sequences. The full test dataset was used to calculate the accuracy of the fine-tuned models. For the in-depth evaluation of attention scores, 1,000 sequences from each group were randomly selected from the test dataset. The same training, test and evaluation datasets were used for DNABERT and Nucleotide Transformer.

#### TATA sequences

To fine-tune the DNABERT and Nucleotide Transformer models for classifying TATA-promoter and non-promoter sequences, we prepared a custom dataset. Real TATA-promoter sequences (300 bp) were extracted from the training and test datasets published in the Nucleotide Transformer paper, which are available on HuggingFace (https://huggingface.co/datasets/InstaDeepAI/nucleotide_transformer_downstream_tasks_revised/tree/main/promoter_tata).

Although the method used in that paper for generating negative control sequences improves model performance, it makes the interpretation of attention scores more difficult. To address this, we created a simpler set of control sequences (300 bp), randomly extracted from non-promoter regions (as defined by the ExPASy EPDnew database), and ensured they did not contain any ‘N’ nucleotides.

The final training dataset consisted of 2,531 TATA-promoter sequences and 2,531 control sequences. The test/evaluation dataset included 106 TATA-promoter and 106 control sequences.

#### Generated fake TATA sequences

To further investigate attention score behavior in an extremely simple task, we created a dataset of randomly generated sequences (300 bp), where the only difference between positive and negative sequences is the presence of the 6-mer “TATAAA” in the positive ones. Both the training and test/evaluation datasets consisted of 2,500 positive and 2,500 negative sequences.

#### Multiple Sclerosis

For scGPT interpretability was investigated across two primary datasets, encompassing both pre-training and fine-tuning stages. The first dataset used to fine-tuned the pre-trained scGPT model was the MS dataset, originally introduced by Schirmer et al.^36^ and obtained from EMBL-EBI (https://www.ebi.ac.uk/gxa/sc/experiments/E-HCAD-35/results). This dataset comprised samples from 9 healthy controls and 12 individuals with MS. During fine-tuning, the model was specifically tailored on a reference partition consisting of healthy human immune cells, with subsequent evaluation focusing on predicting cells afflicted with MS. The reference set was curated to exclude B cells, T cells, and oligodendrocyte Bs and comprised 7,844 cells. The query set contained 13,468 cells. Data processing involved selecting highly variable genes (HVGs), retaining 3,000 genes across both reference and query sets, encompassing 18 distinct cell groups of excitatory neurons, interneurons, oligodendrocytes, glial cells, precursor cells, astrocytes, microglial cells, endothelial cells and phagocytes.

#### Human Pancreas

For the second dataset we used the human pancreas dataset obtained from five scRNA-seq studies by Chen et al. (https://github.com/JackieHanLab/TOSICA)^37^. The five datasets were split into reference and query sets by data sources. The reference set consists of data from two data sources, and the query set contains the other three. Both reference and query sets retained 3,000 genes and ground truth annotations from their original publications. The reference set comprised 10,600 cells spanning 13 distinct cell groups (alpha, beta, ductal, acinar, delta, pancreatic stellate, pancreatic polypeptide, endothelial, macrophage, mast, epsilon, Schwann and T cell), while the query set included 4,218 cells across 11 cell groups (alpha, beta, ductal, pancreatic polypeptide, acinar, delta, pancreatic stellate, endothelial, epsilon, mast and MHC class II).

### Attention score extraction

Scaled dot-product attention is computed as follows:

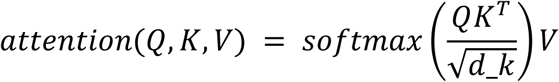

The attention matrix is the result of the softmax on the dot product of the keys and queries after scaling. It is this attention matrix we extract from three representative models in genomics; DNABERT, Nucleotide Transformer, and scGPT, to evaluate the interpretability of attention scores. Specifically, since softmax is applied column-wise, we take the maximum attention score across the row values of the attention matrix in order to extract the maximum attention paid to a specific token.

DNABERT’s attention scores were available for extraction using extract_attentions=True in the setup of the model on Pytorch. They were also available from Nucleotide Transformers HuggingFace implementation as output_attentions=True. To extract attention scores from scGPT, we implemented the model in PyTorch and saved the attention scores matrix in a dictionary within each transformer module such that the attention matrix for every head and every layer was accessible at the end of the prediction for every head and layer.

### Biological annotations

#### DNABERT and Nucleotide Transformer

Given that both DNABERT and Nucleotide Transformer take DNA sequences as input, we used the same strategy to annotate biological features. First, sequences in the evaluation datasets were aligned to the GRCh38 reference genome using PxBLAT^38^ to get their genomic coordinates.

These genomic coordinates were then used to extract individual nucleotide annotations from the biological databases mentioned above. For categorical features, we used the label 0 as “no-feature” and 1 as “feature”. Since both model’s attention calculation operates on “binned base pairs” rather than on individual nucleotides (DNABERT is overlapping k-mers with k=6 and Nucleotide Transformer has non-overlapping k-mers where k=6), binned bp scores were assigned by calculating the mean feature score. GC percentage was directly calculated for each token. Finally, TFs and repeated elements were condensed under families. For each bin, the scores of all TFs belonging to the same family were summed and assigned to that specific family. For repeated elements, we kept the maximum score among all the elements of the same family. All features are listed in Supplementary Table S1.

#### scGPT

The Molecular Signatures Database (MSigDB)^39^ provided the foundation for annotating biological annotations to scGPT’s gene tokens. MSigDB houses a vast collection of gene sets, each an unstructured group of genes linked to specific biological processes (e.g., cell cycle), locations within the genome (e.g., chromosome 1), diseases (e.g., breast cancer), or pathways (e.g., KEGG cell cycle pathway). To enrich scGPT’s gene tokens with biological context, we leveraged the collections: C8 (cell type signatures), C5 (ontology gene sets), and H (hallmark gene sets) for annotation. C8 provides information on cell types by containing gene sets that identify specific cell populations observed in human single-cell sequencing studies. C5 offers a broader functional view by grouping genes based on shared Gene Ontology (GO) annotations (biological process, cellular component, molecular function) or Human Phenotype Ontology (HPO) annotations. H provides a concise representation of core biological processes through “hallmark” gene sets, which are derived by aggregating related gene sets from across MSigDB.

We implemented a filtering step based on gene set representation in our cell populations. For the MS model, gene sets were required to have at least 40 genes expressed in 10% of the samples.

For the Pancreas model, gene sets were required to have at least 35 genes expressed in 10% of samples. Following filtering, each gene token within a cell was assigned a binary value (0 or 1) indicating its membership within a particular gene set. All features are listed in Supplementary Table S1.

**Table 4.** A number of scGPT biological annotations were obtained from MSigDB Hallmark (H), C5 and C8 gene sets.

| Dataset | Total Number of Gene Sets |
| --- | --- |
| Multiple Sclerosis | 81 |
| Human Pancreas | 99 |

### Model and Dataset Overview

**Table 5.**
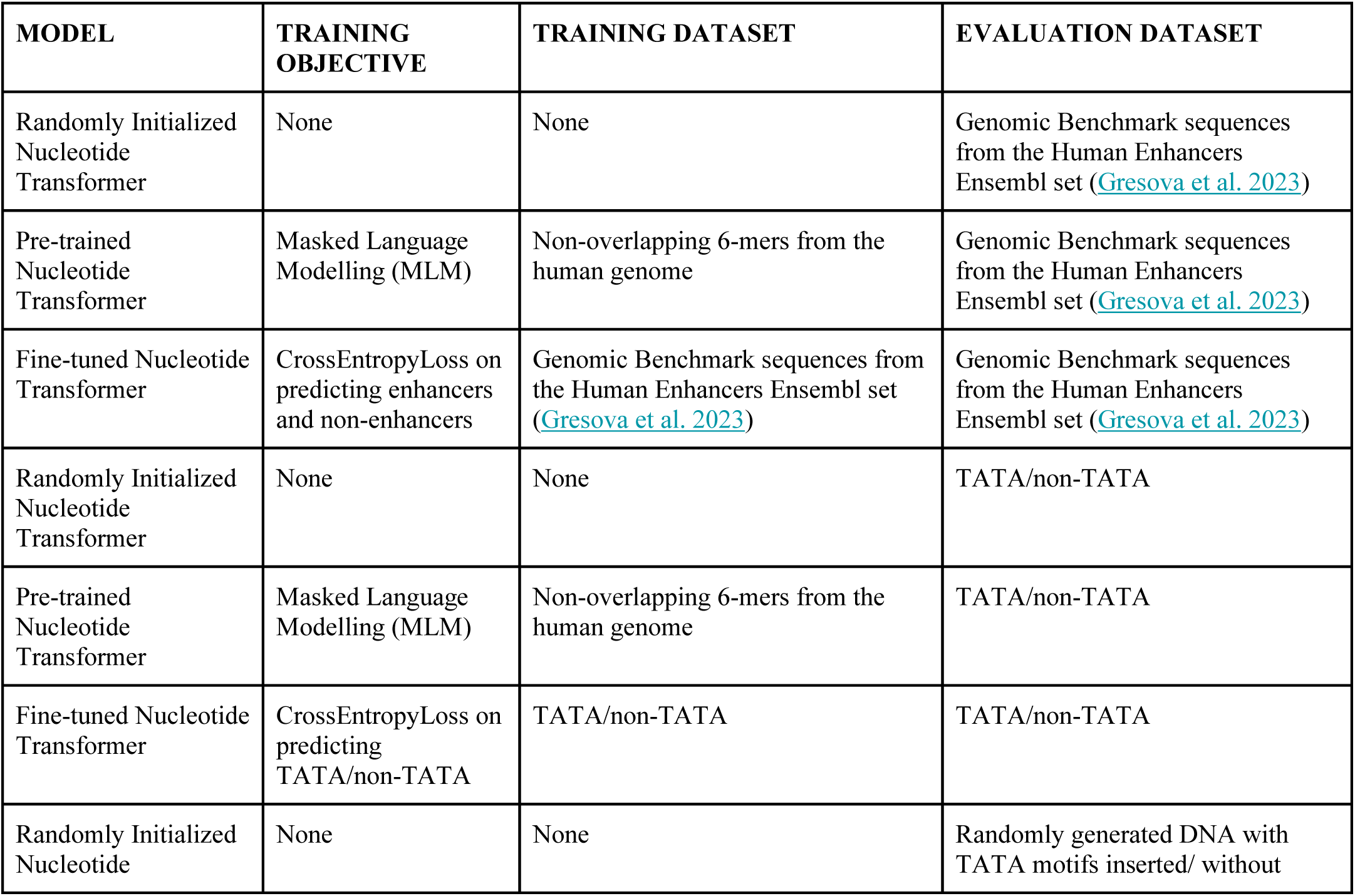
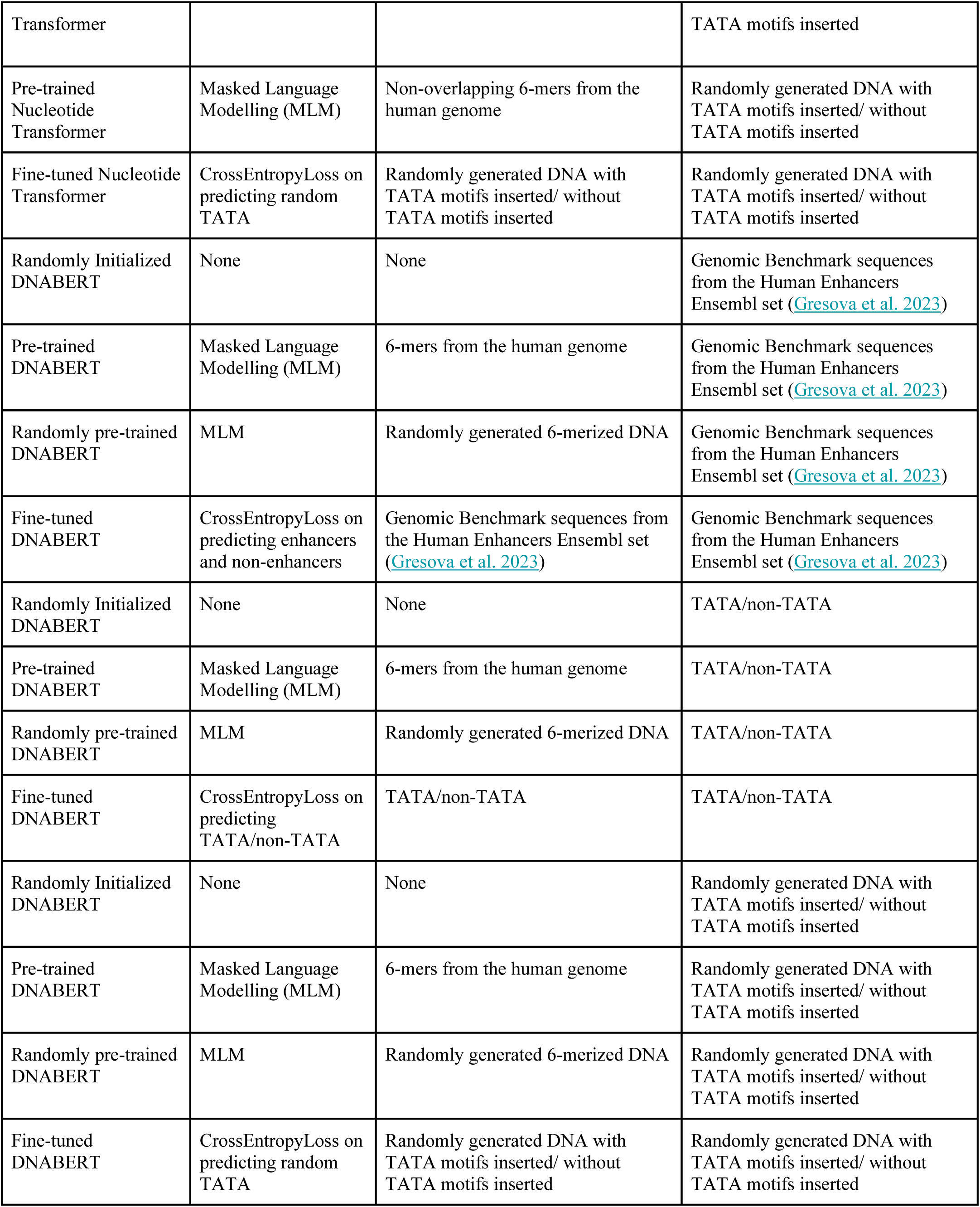
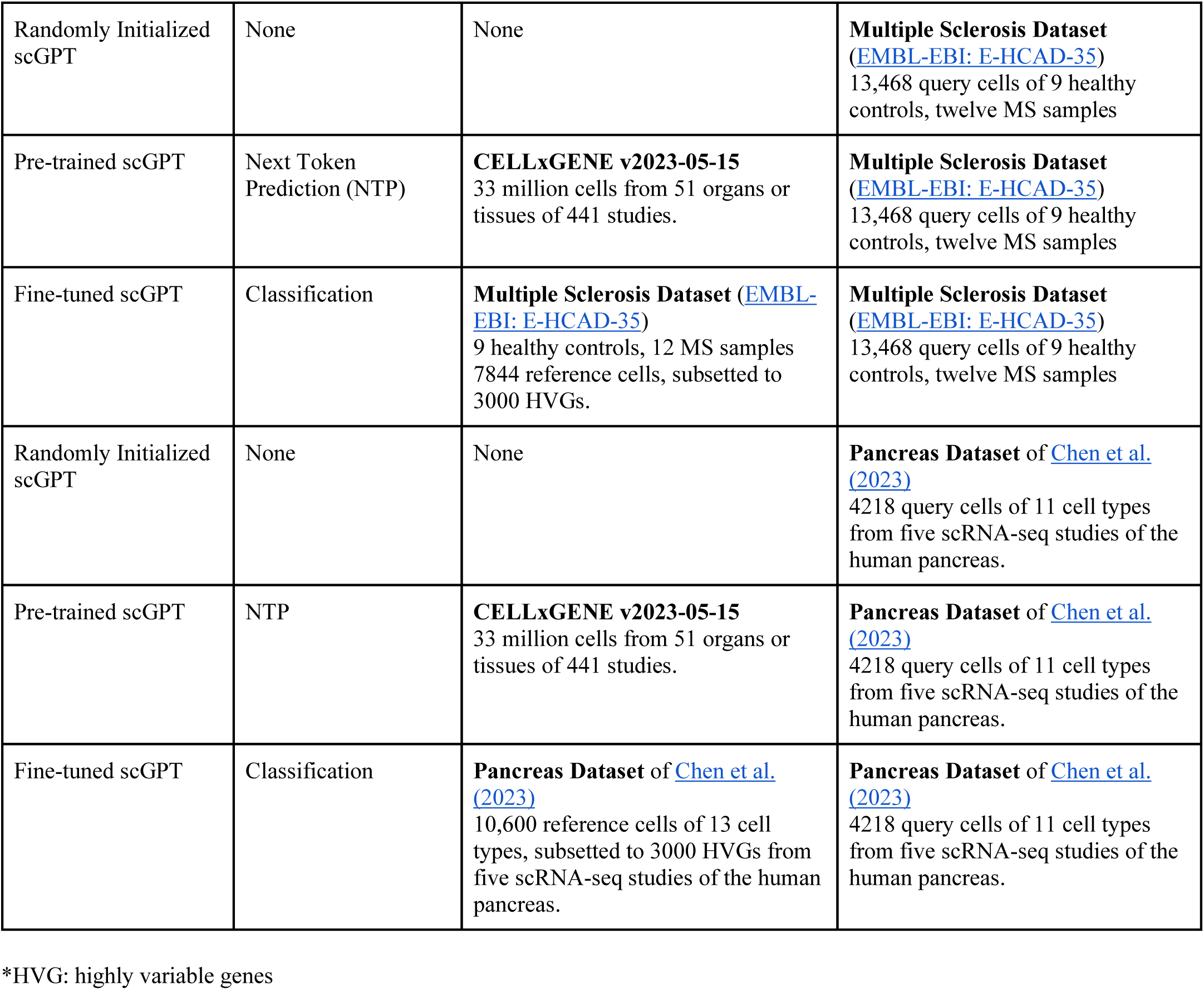
An overview of all the models included in the analysis and their pre-training or fine-tuning dataset, and evaluation dataset (which attention scores were extracted from).

### Spearman Correlations

We employed Spearman rank correlation to analyze relationships between biological annotations and attention scores on tokens within sequences. We used Spearman rank correlations instead of Pearson correlations to capture both linear and monotonic relationships, as our dataset is non-normally distributed and attention scores are unlikely to be linearly associated with biological annotations.

Given the inherent structure of our data, where correlations are performed on sequential tokens, we acknowledge that the assumptions of independence and identically distributed (i.i.d.) samples are not met. The sequential nature of the tokens introduces dependencies, as adjacent tokens are inherently related due to their ordering. Therefore, to assess the significance of our findings, we generated a null hypothesis where attention scores are not related to biological annotation scores.

This is done by generating 100 random shuffles of the attention scores in blocks of 10 tokens, both within and across sequences. By doing this, we preserve some local relationships within attention scores and maintain the overall distribution of both attention and biological annotation scores. Simultaneously, we disrupt the specific associations between attention scores and biological annotations as well as their sequence ordering. After shuffling, we computed Spearman correlation coefficients between the reshuffled attention scores and the biological annotations.

For each biological annotation, we calculated the mean and standard deviation of the correlation coefficients from the shuffled data. We then converted the correlation coefficients into Z-scores by comparing them to the distribution of shuffled coefficients. This method allows us to evaluate whether the observed correlations significantly differ from what might be expected by chance.

Our null hypothesis is that there is no correlation between attention scores and biological annotation presence.

To account for attention head’s activating based on specific context, we conducted analysis subsetted by the labelled inputs set on. For each head we computed attention-score and biological-feature associations using the complete dataset. We also calculated correlations using only the subset of samples from each label, capturing each attention head’s global attention association, and label-specific context associations. This approach allowed us to identify both general features learned by heads and specific features used by specialized heads to predict particular labels.

Features with Z-scores exceeding three standard deviations from the mean (calculated from the 100-shuffled distributions) were considered statistically significant. These significant correlations indicate non-random associations between attention scores and biological annotations, highlighting potential mechanistic links between model attention and underlying biological functions.

### Different tokenization schemes affect the interpretability of models

To validate our method for associating attention heads with biological features, we trained both DNABERT and Nucleotide Transformer on a synthetic TATA dataset. This dataset consisted of 10,000 randomly generated 300 bp sequences (5,000 train, 5,000 test), where positives contained the 6-mer “TATAAA” and negatives did not. Since the TATAAA motif was the only differentiating signal, we expected our method to uncover attention heads associated with it.

Both models revealed heads associated with the TATAAA and ATATAA 6-mers. However, the strongest signals came from heads correlated with GC content. To assess which associations contributed more to task performance, we ablated heads associated with GC or with TATA-related k-mers and measured the impact on classification.

In DNABERT, ablating TATAAA-associated heads led to a larger performance drop than ablating GC-associated heads, as expected (Supplementary Figure S7A<u>-B</u>). But in the Nucleotide Transformer model, removing GC-associated heads had a much greater effect. We hypothesized this was due to its tokenization scheme (non-overlapping 6-mers) which fragments the TATAAA motif into multiple tokens. This likely made the signal harder for heads to learn directly. Instead, GC-related heads may act as indirect detectors by picking up the dip in GC content around the motif.

To test this, we analyzed layer24-head9, a GC-associated head in the Nucleotide Transformer model, and found it showed a marked drop in attention at the TATA site (Supplementary Figure S7C). This suggests that, in this context, GC attention is serving as a proxy for TATA detection. These findings illustrate that tokenization choices can significantly influence how interpretable head-feature associations are using our method.

### Ablation Experiments

We conducted ablation experiments to quantify the functional importance of individual attention heads within genomic transformer models (scGPT, DNABERT, and Nucleotide Transformer).

To isolate and measure each head’s contribution to model performance, we developed a targeted “head zeroing” approach that selectively removes the contribution of specific attention heads while preserving the model’s overall architecture.

Our ablation technique implements a HeadZeroer module that precisely targets the output representation of specific attention heads. For each targeted head, we inserted this module after the attention output projection in the relevant transformer layer. The module operates by identifying the exact hidden state dimensions corresponding to the targeted head (based on head index and dimension size after the multi-head attention concatenation) and setting these values to zero, effectively removing that head’s contribution from downstream layers.

For each model, we measured the performance on the held out test dataset of the baseline fine-tuned model (unmodified model with all attention heads fully functional), the model with mechanistically important heads zeroed out, and a model with layer-matched heads zeroed out.

We evaluated model performance using multiple metrics including classification accuracy and area under the receiver operating characteristic curve (AUC).

The functional importance of ablated heads was quantified by the relative performance decrease compared to the baseline model. By comparing the performance impact of ablating putatively important versus layer-matched heads, we obtained a quantitative measure of our mechanistic understanding of the model’s internal operations. Substantial performance degradation following important head ablation, coupled with minimal impact from layer-matched head ablation, provides strong evidence that our mechanistic interpretations accurately capture functionally relevant aspects of the model’s prediction-making process.

### Interpretability prompting approach

#### OpenAI GPT-4 interpretability prompting

OpenAI recently developed a technique to automate the interpretability of neurons in large language models, such as GPT-2 XL, using GPT-4^21^. They did this using three models, a subject model (GPT-2 XL) they hoped to explain, an explainer model (GPT-4) to generate explanations, and a simulator model (also GPT-4) to provide a way to evaluate these explanations.

Initially, input samples were obtained from particular neurons in the subject model that exhibited the highest levels of activation. Then, the activation of these neurons on specific text/tokens was used to determine the areas of “focus” for these neurons. The goal was to generate researcher-written explanations for real explainable neurons in the model. For example, a neuron that routinely experienced high activation on specific tokens related to Canada could be explained by a researcher as a neuron focusing on Canada-related words (i.e. “Toronto”, “Maple Leafs”, “Canucks”).

Once explanations had been generated for a few real neurons, the authors sent a prompt to the explainer model, GPT-4, that included a few examples of real neurons with real activations and the researcher-written explanations. The prompt concluded with the activations of a new neuron on specific tokens (from one of its highest activating inputs), and asked GPT-4 to explain the new neuron’s behaviour, given the other explained neurons as an example. The explainer model then generated an explanation for the new neuron given the token activation example received.

This explanation was thereafter given to the simulator model (also GPT-4), with a prompt that assumed that the explanation of the explainer model was accurate and comprehensively explained the neuron’s behaviour. Given the explanation of the neuron, the simulator model was asked to generate activations for each token in a particular sequence. That same sequence was then fed into the model to score the activations of the neuron on that sequence. The simulated activations and actual activations were compared, with a higher correlation score indicating a better-explained neuron by GPT-4.

#### Adapted genomic GPT-4 interpretability prompting

Instead of explaining every neuron in these models, we explained every attention head. This was both due to compute limitations, as well as to develop an understanding of specifically how attention heads work in the context of genomic and single-cell transformers.

Unlike in the case of NLP models, the underlying tokens the attention heads in genomic and single-cell transformer models attend to are not interpretable. For an NLP model, higher attention scores on tokens like “Toronto”, “Maple Leafs”, and “Canucks” can be interpreted by humans as Canada-related, whereas higher attention scores on tokens with higher GC content are not immediately obvious. As such, we developed annotations for input sequences fed into the model such that every token was scored based on a subset of biological annotations of interest.

We then ran Spearman rank correlations for every head and layer to correlate the attention scores on each token with the biological annotations of interest. The coefficients of these models were used similarly to the raw activation scores in OpenAI’s method. While OpenAI scaled attention scores for every token between 0-10, we did not scale the coefficients of each biological annotation. As the biological annotations are interpretable to expert humans, the same way human-language tokens are, we sent the biological annotations and coefficients to GPT-4 to explain, leveraging its apparent biological knowledge base, using a very similar format to OpenAI.

Unfortunately, we could not use GPT-4 as a simulator model in the same way OpenAI did, as we would not be predicting attention scores on tokens per sample, but rather coefficients of Spearman rank correlations of the attention scores for each biological annotation across a dataset.

#### Prompting

Mimicking the OpenAI researcher-written explanation style, we prepared our own explanation example for GPT-4 prompting. In this sense, in a short paragraph, we explained to GPT-4 the experiment we were performing and what we expected from it, detailing the input provided. For example, for DNABERT we used the following prompt: *“Analyze this attention head from a DNA transformer model fine-tuned to predict enhancer vs non-enhancer sequences*.

*Format:*

*Feature name: [Feature coefficient for enhancer, non-enhancer]*

*Higher absolute values = stronger association*

*Positive = positive association, Negative = negative association*

*Instructions:*

1. *Identify the most informative features (those with the largest coefficient magnitudes)*.
2. *Note how feature importance differs between enhancer and non-enhancer sequences (e.g. if |enhancer coefficient| > |non-enhancer coefficient|, that feature is more relevant in enhancer sequences)*.
3. *Apply your biological knowledge to provide a mechanistically meaningful and biologically accurate explanation*.

*Output ONLY the following format:*

*‘Head Name: [3–5 word mechanistically descriptive name] Explanation: [1–2 plain, specific sentences describing what this head captures biologically. Refer to specific features and how they differ between enhancer and non-enhancer sequences.]’*

*Failsafes:*

*If the head does not meaningfully distinguish enhancer from non-enhancer, say so directly. If the observed associations contradict established biology, do not try to rationalize or justify them—simply state that the pattern is unclear or inconsistent*.

*Avoid vague phrases like “associated with” or “element sensitivity.” Be direct and specific*. *Do **not** include any bullet points, summaries, or explanations outside the requested format.”*. For prompting we used GPT-4.1 (gpt-4.1-mini-2025-04-14) API. The average cost of prompting head explanations for one attention head was $0.01 USD (Making summarizing the Nucleotide Transformer model with 464 heads cost a total of $4.64).

GPT-4 generated explanations for the heads can be found in the genome-head-interpreter/preprocessing/data/explanations/ section of the code.

### DNABERT and Nucleotide Transformer k-mer Analysis

#### Head analysis by label

We examined which k-mers and attention heads were most relevant to the classification task. First, tokens were grouped according to their sequence label –either belonging to an enhancer or TATA-promoter sequence, or to a control sequence. To minimize bias due to sample size differences, we excluded k-mers that appeared fewer than 15 times in DNABERT or fewer than 3 times in Nucleotide Transformer. Next, we computed i) the global mean attention score of each head for each group, considering the attention scores of all k-mers, and ii) the mean attention score of each individual k-mer for each head. For each label, we then calculated the difference between the individual k-mer attention score and the global mean head score. This allowed us to identify k-mers that showed attention patterns distinct from the overall head behavior across conditions.

## Data availability

The datasets and their labels are available at Zenodo: https://zenodo.org/records/15484539.

## Code availability

The scripts used to extract the attention scores, process the results and generate the figures shown in the paper can be found on GitHub: https://github.com/meconsens/interpretability.git. Raw attention scores for each of the models are stored in Zenodo: https://zenodo.org/records/15484539.

## Supplementary

**Supplementary Figure S1.** Scatter plots showing the difference between the mean head attention score and the specific k-mer mean attention score for A) DNABERT and B) Nucleotide Transformer models fine-tuned on the three different datasets: TATA, fake TATA and enhancers. Positive controls refer to real TATA promoter or enhancer sequences, while negative controls are sequences not labeled as TATA promoters or enhancers. k-mer heads with scores beyond three standard deviations from the diagonal are highlighted in red, and the top three k-mer heads are labeled.

**Supplementary Figure S2.** Z-scores for each feature and attention head comparing pre-trained and randomly initialized models in A) DNABERT, B) Nucleotide Transformer, and C) scGPT, across the different datasets. k-mer heads with scores beyond three standard deviations from the diagonal are highlighted in red, and the top three k-mer heads are labeled. The highest activations appear along the y-axis, indicating that some features are learned during pre-training.

**Supplementary Figure S3.** DNABERT z-scores for each feature and attention head comparing random pre-trained models with A) random initialized, B) pre-trained and C) fine-tuned models, across the TATA promoters, fake TATA and enhancers datasets. Positive controls refer to real TATA promoter or enhancer sequences, while negative controls are sequences not labeled as TATA promoters or enhancers. k-mer heads with scores beyond three standard deviations from the diagonal are highlighted in red, and the top three k-mer heads are labeled. The pattern observed in A), comparing randomly pre-trained with randomly initialized models, is similar to that in Supplementary Figure S2 (pre-trained vs. randomly initialized), suggesting that random pre-training may capture some generalizable biological signals.

**Supplementary Figure S4.** Z-scores for each feature and attention head comparing pre-trained and fine-tuned models in A) DNABERT, B) Nucleotide Transformer, and C) scGPT, across the different datasets. k-mer heads with scores beyond three standard deviations from the diagonal are highlighted in red, representing features and heads whose attention changes during fine-tuning. Top three k-mer heads are labeled.

**The Supplementary Figure S5.** Additional ROC curves complementing those in Figure 4, showing results from further ablation experiments. A) DNABERT and Nucleotide Transformer models fine-tuned on the enhancer dataset, with ablation of heads correlated with the GC content feature. B) scGPT fine-tuned on the Multiple Sclerosis and Pancreas datasets, with ablation of heads correlated with the “Synapse” cellular component GO term and the “Pancreas ductal cell” label, respectively. The dotted diagonal line represents a random classifier (AUC = 0.5).

**Supplementary Figure S6.** DNABERT ablation experiments targeting heads correlated with the TATAAA k-mer: A) regardless of correlation direction, B) positively correlated heads, and C) negatively correlated heads. Heads negatively correlated with the TATAAA k-mer appear to be more important for model performance, as ablating just 5% of them leads to a noticeable drop in accuracy compared to positively correlated heads. The dotted diagonal line represents a random classifier (AUC = 0.5).

**Supplementary Figure S7.** ROC curves for the head ablation experiments on the fake TATA dataset for A) DNABERT and B) Nucleotide Transformer models. For both models, we show the effect of ablating heads correlated with either the GC content or the TATAAA k-mer feature. The dotted diagonal line represents a random classifier (AUC = 0.5). C) Mean attention scores and mean GC content (%) across 100 bp surrounding the TATAAA k-mer for head 9 in layer 24 (identified as a GC feature head) of the Nucleotide Transformer in the pre-trained, fine-tuned, and randomly initialized models. The drop in GC content at the TATAAA k-mer corresponds to a peak in attention scores.

## Contributions

M.E.C.: Conceptualization (supporting), data curation (equal), formal analysis (equal), methodology (equal), project administration (equal), software (lead), visualization (equal), writing – original draft preparation (lead), writing – review & editing (equal)

A.D.-N.: Data curation (equal), formal analysis (equal), methodology (equal), validation (equal), visualization (equal), writing – original draft preparation (equal), writing – review & editing (equal)

V.C.: Data curation (equal), formal analysis (equal), methodology (equal), software (equal), validation (equal), visualization (supporting), writing – original draft preparation (equal), writing – review & editing (equal)

L.S.: Supervision (supporting), resources (supporting), writing – review & editing (equal)

H.H.H.: Supervision (supporting), writing – review & editing (equal)

A.M.: Supervision (supporting), writing – review & editing (equal)

B.W.: Conceptualization (lead), funding acquisition (lead), project administration (equal), supervision (lead), resources (lead), writing – review & editing (equal)

## Acknowledgements

Thanks to the developers and contributors of DNABERT, Nucleotide Transformer, and scGPT models. Appreciation goes to OpenAI for providing GPT-4, which plays a crucial role in our interpretability analysis.

## Funding

M.E.C. was supported by the Schwartz Reisman Institute for Technology and Society at the University of Toronto through a Schwartz Reisman Fellowship, and by the Natural Sciences and Engineering Research Council of Canada (NSERC). V.C. was supported by the Princess Margaret Cancer Centre through the Cancer Digital Intelligence Spark Award. D-N.A was supported by the Ontario Genomics-CANSSI Ontario Postdoctoral Fellowship program in Genome Data Science.

## Notes

https://zenodo.org/records/15484539

### Competing Interest Statement

B.W. is currently employed as the Senior Vice President and Head of Biomedical AI at Xaira Therapeutics. He also serves as a scientific advisor to Deep Genomics, Shift Biosciences, and VieCure Inc.

